# Maintenance of memory by negative-feedback of synaptic protein elimination: Modeling KIBRA-PKMζ dynamics in LTP

**DOI:** 10.1101/2024.09.25.614943

**Authors:** Harel Z. Shouval, Changchi Hsieh, Rafael E. Flores-Obando, David Cano, Tara E. Tracy, Todd C. Sacktor

**Affiliations:** Department of Neurobiology and Anatomy, University of Texas Medical School, Houston, TX 77030, USA; Department of Electrical and Computer Engineering, Rice University, Houston, TX 77005, USA; Department of Physiology and Pharmacology, State University of New York Downstate Health Sciences University, Brooklyn, NY, 11203. USA; Department of Neurology and Anesthesiology, State University of New York Downstate Health Sciences University, Brooklyn, NY, 11203. USA; School of Medicine, State University of New York Downstate Health Sciences University, Brooklyn, NY, 11203. USA; Buck Institute for Research on Aging, Novato, California, 94945, USA

## Abstract

Activity-dependent modifications of synaptic efficacies are a cellular substrate of learning and memory. Current theories propose that the long-term maintenance of synaptic efficacies and memory is accomplished via a positive-feedback loop at the level of production of a protein species or a protein state. Here we propose a qualitatively different theoretical framework based on negative feedback at the level of protein elimination. This theory is motivated by recent experimental findings regarding the binding of *PKMζ* and KI-BRA, two synaptic proteins involved in maintenance of memory, and on how this binding downregulates the proteins’ degradation. We demonstrate this theoretical framework with two different models. First, a simple abstract model to explore generic features of the negative-feedback process. Second, a biophysical model based on *PKMζ*-KIBRA dimers that cooperatively form larger complexes at active synapses. These larger complexes have slower degradation and diffusion, allowing for bistability of potentiated and unpotentiated synaptic states. The results of these models are qualitatively consistent with existing experiments showing reversal of long-term potentiation and erasure of long-term memory by inhibition of KIBRA-*PKMζ* interactions. The theory generates novel predictions that could be experimentally tested to further validate or reject the negative-feedback theory.

## 1 Introduction

We all have memories that date back to our youth; we remember the house we lived in at age four; we remember a favorite or least favorite schoolteacher. These memories are believed to be encoded primarily via long-lasting synaptic plasticity that persistently modifies specific neuronal circuits in our brain (Martin et al., 2000; Whitlock et al., 2006; Yang et al., 2016; Langille and Brown, 2018), and at the molecular level this is achieved by altering the numbers or conformational states of particular proteins in these synapses (Bear and Malenka, 1994; Nicoll, 2017). However, this putative cellular basis of memory relies on proteins that typically have lifetimes orders of magnitude shorter than the memory. Here exactly lies a fundamental problem of long-term memory and synaptic plasticity: How can memories be stored for a human lifetime on the basis of proteins that are continuously degrading (Crick, 1984; Lisman, 1985). Moreover, since synaptic plasticity is a synaptic rather than a whole cell event, what affects the lifetimes of individual synaptic efficacies and hence memory is likely not the lifetime of a protein in the cell, but the time that a protein typically dwells in a synapse. This variable, the synaptic lifetime, may well be significantly shorter than the protein lifetime due to diffusion and trafficking of proteins.

To account for how such an inherent instability of memory can be overcome, different theories have been proposed (Lisman and Zhabotinsky, 2001; Bhalla and Iyengar, 1999; Aslam and Shouval, 2012; Jalil et al., 2015). Most theoretical models assume a common mathematical structure based on a positive-feedback mechanism that can generate bi-or multi-stability. The biophysical instantiation of the feedback varies from model to model. This feedback could be based on single species feedback loops such as autophosphorylation (Lisman, 1985; Lisman and Zhabotinsky, 2001), or more complex multi-molecule post-translational feedback loops (Bhalla and Iyengar, 1999), or, alternatively, on positive-feedback loops operating at the level of the regulation of translation (Aslam et al., 2009; Jalil et al., 2015; Richter and Klann, 2009; Ogasawara and Kawato, 2010). Note that despite the different biophysical substrates of these theories, their mathematical structure is similar.

One prominent candidate for a molecule that is necessary for the maintenance of late-phase long-term synaptic plasticity (L-LTP) and long-term memory is the constitutively active form of atypical protein kinase C, *PKMζ*(Sacktor et al., 1993; Ling et al., 2002; Pastalkova et al., 2006). The levels of *PKMζ* persistently increase after L-LTP, in selective postsynaptic compartments of memory encoding neurons for at leas a month in hippocampus (Hsieh et al., 2021) and neocortex (Gao et al., 2018). Blocking *PKMζ* activity reverses L-LTP and established memory (Pastalkova et al., 2006; Tsokas et al., 2024). How this persistent increase is maintained despite protein turnover is still unclear. Previous models (Ogasawara and Kawato, 2010; Jalil et al., 2015) have postulated that this might occur due to a feedback loop based on enhanced translation in synaptic spines. The results of these models seem consistent with experiments; however their fundamental assumption and the mechanism for such persistently increased translation have not been elucidated (though see (Westmark et al., 2010)).

Recent results have shown that maintenance of synaptic plasticity and memory depends on the binding between *PKMζ* to another postsynaptic molecule, KIdney BRAin protein (KIBRA) (Hu et al., 2017; Ferguson et al., 2019; Tsokas et al., 2024). During L-LTP the synaptic concentration of *PKMζ* bound to KIBRA increases selectively in activated pathways (Tsokas et al., 2024). Agents that specifically interfere with the binding of *PKMζ* and KIBRA can reverse established L-LTP and long-term memory as late as a month after memory formation (Tsokas et al., 2024). In contrast, these inhibitors do not affect baseline synaptic transmission and do not affect the binding of other kinases such as other atypical protein kinase C molecules and *Ca*^2+^/calmodulin-dependent protein kinase, II (CaMKII) to KIBRA (Tsokas et al., 2024). In addition, previous results show that the binding of *PKMζ* to KIBRA shields *PKMζ* from proteasomal degradation (Vogt-Eisele et al., 2014). Together, these results suggest that the increase of *PKMζ* magnitude in synapses during long-term plasticity and memory arises from a decrease in protein turnover, because KIBRA is a postsynaptic scaffold acting as a synaptic tag for *PKMζ*, and possibly decreased diffusion rather than from an increase in synthesis. There are currently no mathematical theories of memory maintenance based on the conditional decrease of protein elimination.

Motivated by these results we propose a novel theory based on negative feedback of protein elimination. In the context of this general theoretical framework, the concept of elimination incorporates several molecular processes, including classical protein degradation, but also changes of the conformational state of a protein from an active to an inactive state, as well as the diffusion or trafficking of proteins out of the synaptic compartment. Here we explore two specific instantiations of the negative-feedback theory, a simple single species model to explore the principles of maintenance by negative feedback and a more complex biophysical model motivated by the binding of *PKMζ* and KIBRA (Vogt-Eisele et al., 2014; Hu et al., 2017; Ferguson et al., 2019; Tsokas et al., 2024). Using these models we will outline the general properties of the negative-feedback theory, examine under what general conditions it can produce stability, and propose experimentally testable predictions.

## 2 Results

This paper aims to explain how maintenance can be achieved by negative feedback of protein elimination. Before turning to the analysis of the negative-feedback theory, we will begin by examining a simplified synaptic state model (section 2.1), and explain how positive-feedback theories operate in this framework (section 2.2). This section does not describe novel work but is a simple generic description of most previous models, attempting to explain maintenance of synaptic efficacies. We provide this section in order to contrast these previous theories with the negative-feedback framework. Using the same 1D dynamical structure we will develop a simple instantiation of the negative-feedback theory (section 2.3). We will show that this model can result in bistability and even generate a continuous attractor. We also outline some generic testable predictions of the theory. Finally in section 2.4, we describe a biophysical model to account for the negative feedback of degradation arising from interactions between *PKMζ* and KIBRA, as observed experimentally (Vogt-Eisele et al., 2014; Tsokas et al., 2024). This model is not a 1D model, as it is described by 9 coupled dynamical equations, but it generates a similar qualitative behavior as the simpler 1D model of negative feedback. The biophysical model suggests a possible mechanistic basis for the origin of the negative feedback and it allows us to simulate the effect of inhibiting the binding of *PKMζ* and KIBRA, which has been shown experimentally to reverse L-LTP and disrupt long-term memory maintenance (Tsokas et al., 2024).

### 2.1 The general mathematical structure of long-term synaptic plasticity in 1D

Synaptic plasticity is a complex high-dimensional process, but here for simplicity we assume a simple 1D model. In some cases the dimensionality of such high-dimensional dynamical systems can be rigorously reduced (Aslam and Shouval, 2012; Agarwal et al., 2012; Gabbiani and Cox, 2017); here we simply assume this can be done. In this 1D system the variable *P* represents the species of interest. This species could be a concentration of a protein, the concentration of a specific state of a protein, for example phosphorylated protein, or it can even represent a complex of bound proteins. In terms of this 1D system a dynamical equation that governs the concentration of this species (P) has the form:

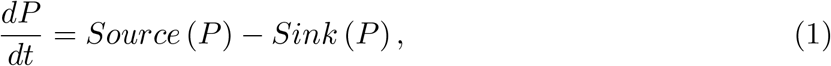

where the source (production) term includes everything that brings *P* into the compartment, including diffusion, trafficking, protein synthesis, and post-translational modifications such as phosphorylation, which produce this form of the protein within this compartment, or the binding reaction of different proteins to a compound. The sink term includes every mechanism that eliminates P from the compartment, including diffusion, protein turnover, and processes such as dephosphorylation or unbinding. If dimensionality reduction techniques are used, the variable *P* could also be some transformed variable representing a combination of some other elementary variables. Note that both the sink and source terms depend on *P*, itself, as both the positive-feedback and the negative-feedback theories require different forms of this dependence on *P*.

Maintenance theories are typically based on equations that have several stable fixed points, and these stable fixed points represent stable states of synaptic efficacy that persist despite ongoing protein degradation and diffusion. In order to have several fixed points, the sink and source terms must cross several times (Fig. 1A), and for this to happen at least one of these two terms must be nonlinear. Most current theories, which depend on positive feedback assume that the interesting non-linear part resides in the source term, and that the sink term is linear. The non-linearity of the source term is assumed to arise from processes such as autophosphorylation or the proteins’ control of its own translation rate within the compartment. In the negative-feedback framework, the interesting nonlinear term is in the sink term, and will be due to negative feedback of one or more of the processes involved in the elimination of proteins from the compartment, which include protein degradation, diffusion, and trafficking.

**Figure 1.**
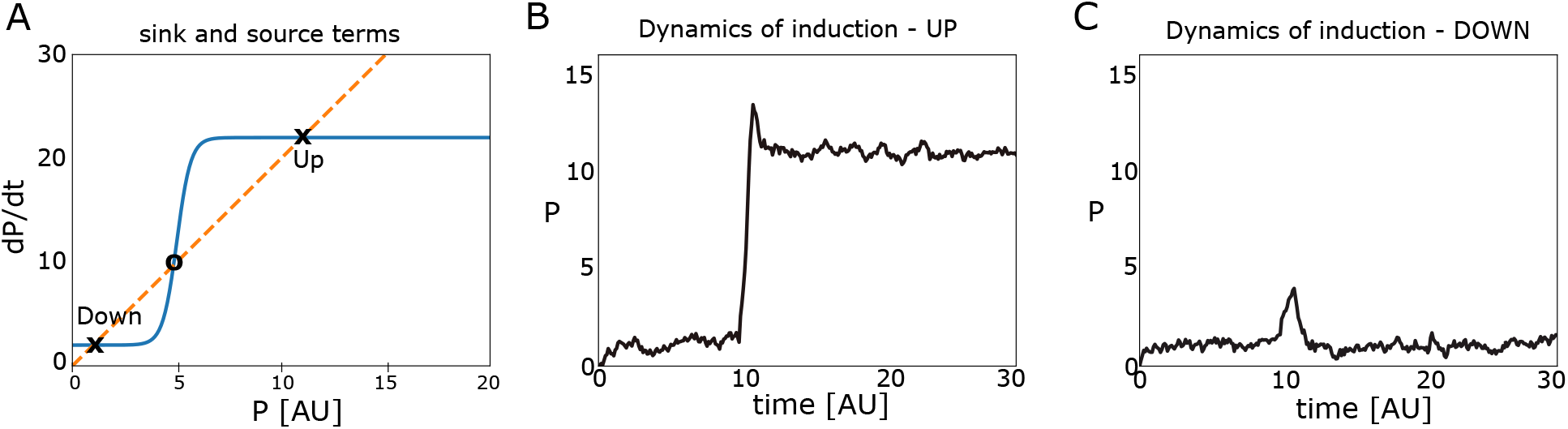
Bistability with positive feedback in the source term. **A.** Sink term (dashed red line) and source term (blue line). These lines cross at three points, two of these are stable fixed-points (x) and one is unstable (o). **B**. Dynamics of L-LTP induction, triggered by a large pulse of a current of P starting at time T=10 [AU]. Here, the system successfully shift from the DOWN to the UP state (dynamics include noise). **C**. With a smaller input pulse there is only a transient change and the system returns to the DOWN state.

### 2.2 Positive feedback theories of maintenance

In positive-feedback theories, the source term is more complex and is the mechanistic origin of synaptic stability, while the sink term is usually assumed to be linear of the form: *Sink* (*P*) = *λP*, where *λ* is the elimination rate constant. More complex, yet monotonic sink terms typically do not alter the qualitative behavior of the system. When the source term has a nonlinear shape, for example, like the source term in Figure 1A, then the sink and source terms cross several times. When the source and sink terms cross, the left hand side of equation 1 is zero, and hence *dP/dt* = 0, which is the definition of a fixed point. In the example shown in figure 1 they cross 3 times, two of these crossings denoted by UP and DOWN are stable fixed points, which correspond to synaptic efficacy in its basal state (DOWN) and after L-LTP (UP). The intermediate state is not stable which means that if the synapse is in that state, small fluctuations will drive the system away from that state towards either the DOWN or UP states.

In Figure 1B we show the simulated induction of LTP in such a system, using a short duration of an additional input current of P. If this current is sufficient, L-LTP is induced and maintained (Fig. 1B). If the current is too small, there is a transient increase in P but there is no maintenance (Fig. 1C). These simulations include a noise term to show that the stable fixed-points are indeed preserved when noise is added (Fig. 1B,C). Noise here was added phenomenologically, and not using a rigorous single-molecule approach (Gillespie, 2002; Agarwal et al., 2012). This approach to adding noise, which is used to understand the qualitative rather than quantitative effects of noise, is used throughout this paper. Note that it is essential that the non-linearity of the source term be steep enough, or ultrasensitive to obtain bistability. If the source term is not ultrasensitive, then the system has only a single fixed point.

The origin of the non-linearity (Fig. 1A) in such systems can arise from various types of molecular interactions, for example auto-phosphorylation (Lisman, 1985), or the up regulation of the translation of new protein in a protein-dependent manner (Aslam and Shouval, 2012; Jalil et al., 2015). These different molecular and cellular mechanisms are mathematically similar as they are all based on positive feedback. The mathematical formulation of these different biophysical processes should be described by additional variables, and therefore, the complete theory is faithfully characterized by a higher-dimensional system. However, within the crucial dimension described by equation 1 the implication of the positive feedback is a steep non-linearity. As shown, positive feedback theories can be bistable, however, whether such systems indeed manifest bistability depends on the other variables in the original higher dimensional system, and on the systems parameters.

For one simple example of a positive-feedback theory, proposed by Lisman (Lisman, 1985), it is possible to reduce the higher dimensional system to a 1D system, and exactly solve for the fixed-points of the system. This was done approximately in Lisman’s original 1985 paper, and we subsequently derived a precise reduction to a 1D equation (Agarwal et al., 2012). This is one example that demonstrates how it is analytically possible to reduce a higher dimensional system to a 1D system, in which the non-linear interactions can generate bistability. More complex and realistic models have been developed using the autophosphorylation of CaMKII and the details of the CaMKII holoenzyme (Lisman and Zhabotinsky, 2001; Miller et al., 2005). Although such models are much more complex, their steady states can sometimes be calculated analytically (Gabbiani and Cox, 2017), and their qualitative behavior may be captured by the 1D model. Another source of a positive-feedback loop can arise from the regulation of translation within synaptic spines, and previous models have modeled the regulation of the synthesis of CaMKII via the protein CPEB1 (Aslam et al., 2009; Aslam and Shouval, 2012) or the regulation of the synthesis of *PKMζ* (Jalil et al., 2015). Although the biological mechanisms are totally different, these different models share a similar mathematical structure.

Positive-feedback models can also generate multi-stability as we have previously shown (Jalil et al., 2015) if the source term has a more complex shape due to multiple feedback mechanisms. Such multi-stability has been both modeled and synthetically constructed in whole mammalian cells at the level of transcription (Zhu et al., 2022).

### 2.3 Stability due to negative feedback of protein turnover: abstract model

The negative-feedback theory requires a non-linear sink term and for simplicity one might assume a constant source term. Here we explore a very simple instantiation of this theory in which we will assume a single protein *P* with a variable sink term and a single synaptic compartment. We will explore below a more complex model (section 2.4), but first we will present a very simple model to illustrate the concept.

Assume for now, that there is a single protein that can undergo a steep transition between two states. In one of these states the degradation rate coefficient (*λ*_1_) is fast; in the other state the degradation rate coefficient (*λ*_2_) is slower (*λ*_1_ *> λ*_2_). The variables *P*_1_ and *P*_2_ denote the concentration of protein in states 1 and 2, respectively, and the total protein concentration is *P* = *P*_1_ +*P*_2_. We define a state function *f*(*P*, Θ_*P*_) that determines the fraction of protein in state 2, and assume that it depends on the total concentration of the protein *P*, and a set of parameters Θ_*P*_. According to these definitions, *P*_2_ = *f* (*P, θ*_*P*_)*·P*, and *P*_1_ = (1 — *f* (*P, θ*_*P*_)) *· P*. Under these assumptions the dynamical equation would take the form:

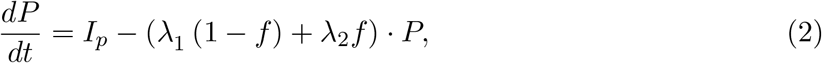

where *I*_*P*_ is the source term, which for simplicity we assume is constant except during the induction phase, in which it is transiently increased. We have omitted the arguments of the state function *f*, for simplicity. The function *f* should be itself modeled as a dynamical process as well, but let us assume we can ignore these dynamics (maybe they are much faster) and that they could be replaced by a function that accounts for their steady-state behavior. For such a 1D system, fixed-points are formed when the source term crosses the sink term. In Figure 2A-C we present these two curves for several different choices of *f*. We use a sigmoidal state function of the form *f*(*x, β, θ*) = 1*/* (1 + *exp*(—*β*(*x* — *θ*))) with two parameters, a slope denoted as *β* and half-maximum denoted as *θ*. In the limit *β* →∞ the sigmoid becomes a step function.

**Figure 2.**
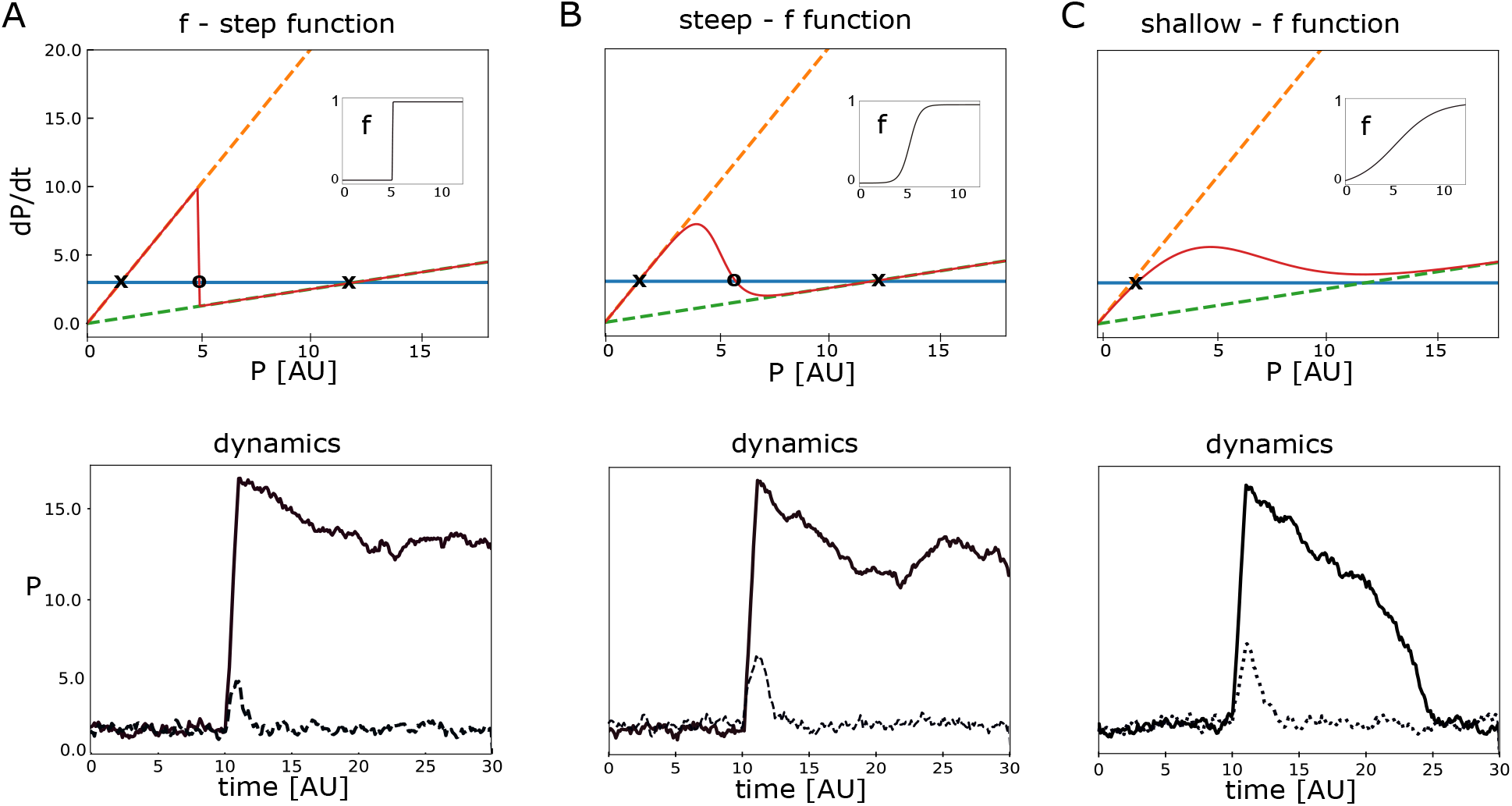
Bistability of a negative feedback model. **A.** Bistability in the case of a state function (f) being a step-function (*β* → ∞, *θ* = 5). Above - A constant source (blue line) term, and two different sink terms with different values of *λ* (dashed orange and green lines). The resulting effective sink term (from eq. 2) given a step state function f (red line). The shape of the state function used is shown in the inset. The effective sink term crosses the source term at 3 points. Stable fixed-points are marked with **x**; unstable fixed-points with **o**. Below - simulations of the induction of L-LTP with an additional noise term. Solid line - large initial stimulus, Dashed line-sub-threshold initial stimulus. **B**. Bistability with f a steep sigmoid function (*β* = 2, *θ* = 5). Top-The resulting effective sink (eq. 2) term given a steep sigmoidal state function f. The shape function used is shown in the inset. Below - as in A, for the steep sigmoid. **C**. Mono-stability is obtained when f is not steep enough (*β* = 0.5, *θ* = 5). Above - shows source and effective sink terms, as in A and B. There is only one intersection between the source and sink terms. Below-simulations always converge to the same fixed point; the system is mono-stable.

The first two examples, in which the state-function is a step-function (Fig. 2A) and a steep sigmoid (Fig. 2B), exhibit bistability. When the state function is not sufficiently steep (Fig. 2C), the system has only a single stable fixed-point. The lower subplots in Figure 2 show simulations in which we emulate the induction of L-LTP with a transient increase in protein synthesis (*I*_*P*_). When the system is bistable L-LTP can be induced (Fig. 2A,B - below); when it is monostable only a transient increase in P is obtained (Fig 2C - below). These simulations include a noise term, responsible for the fluctuations observed in the dynamics. With the given magnitude, these fluctuations, do not destabilize the fixed-points; demonstrating that they are indeed stable. In this bistable system there are two stable fixed points denoted as the DOWN and UP state. We denote the total protein levels in the DOWN and UP states as *P* ^*D*^,*P* ^*U*^ respectively. The magnitude of the two states of the protein in the DOWN and UP fixed points are denoted as: 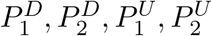, respectively. The total protein concentration is each state is 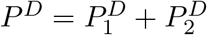 and 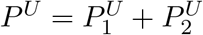.

In the limiting case when *f* is a step function (*β* → ∞), there is a simple relationship between the levels of the total protein in the UP and DOWN states and the degradation rate coefficient, of the form:

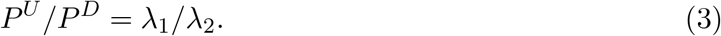

This relationship is also approximately true for a large range of *f*—function parameters that are steep enough to maintain bistability, and can therefore be seen as a prediction of the model. Subsequently we will test if this prediction generalizes to more complex realizations of the theory.

A necessary condition for obtaining bistability in this model is that the total amount of *P*_1_ in the UP state 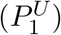 must be lower than in the DOWN state 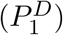. red In the negative-feedback model: 1) the production of protein formed must equal the turnover of protein for any stable state, and 2) the total turnover rate coefficient in the UP state must be lower than in the DOWN state. In the model, the production of protein by new synthesis is constant except during induction. As a consequence, the level of the different forms of the proteins in the two states is entirely controlled by differences in turnover. The turnover coefficients of the *P*_1_ and *P*_2_ forms of the proteins also do not change in the UP and DOWN states. Therefore, as *P*_2_ increases in the UP state, the only way to achieve an equal amount of total protein turning over in the DOWN and UP states is for the amount of *P*_1_ in the UP state to decrease.

The almost all-or-non scenario shown in Figure 2 is not essential for bistability. Consider a different *f* function that saturates at a value *f*_*max*_ *<* 1 for example: *f*(*x, β, θ*) = *f*_*max*_*/* (1 + *exp*(—*β*(*x* — *θ*))). We will now assume that in the DOWN state the value of *f* ≈ 0 and in the UP state it is *f* ≈ *f*_*max*_; an assumption that is approximately correct for a sharp *f* function. In such a case the protein level in the DOWN state is still approximately 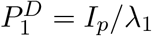, and 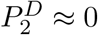. In the UP state the concentrations are approximately:

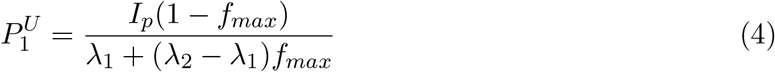

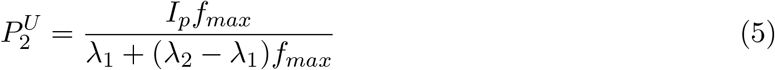

In order to obtain stability under such conditions, it is also useful to have a much lower degradation of the *P*_2_ protein, that is, *λ*_2_ *<< λ*_1_. Using this condition, we find that:

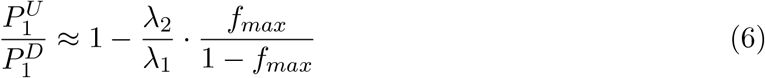

This implies that if indeed *λ*_2_ *<< λ*_1_, the total amount of *P*_1_ in the UP state could be quite close to its amount in the DOWN state, although still necessarily lower. An example of this can be seen in Figure 3. Here, the ratio *λ*_1_*/λ*_2_ = 100, and *f*_*max*_ = 0.85. According to the theoretical approximation above (eq: 6) this yields a ratio of 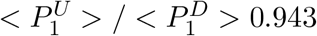. Compare this the simulations, where 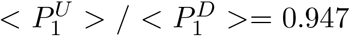 (Fig. 3), when averaged over the periods in each different state. As in the all-or-non case, the condition that the level of quickly degrading proteins is lower in the UP state than in the DOWN state is a necessary condition for bistability. In Figure 3B we also show the total concentration of the protein *P*_*total*_ = *P*_1_ + *P*_1_. The level of *P*_*total*_ is shown by the black line, in the DOWN state *P*_*total*_ = *P*_1_ and the two curves overlap. In the UP state *P*_*total*_ is a little larger than 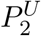 because in the scenario 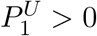.

**Figure 3.**
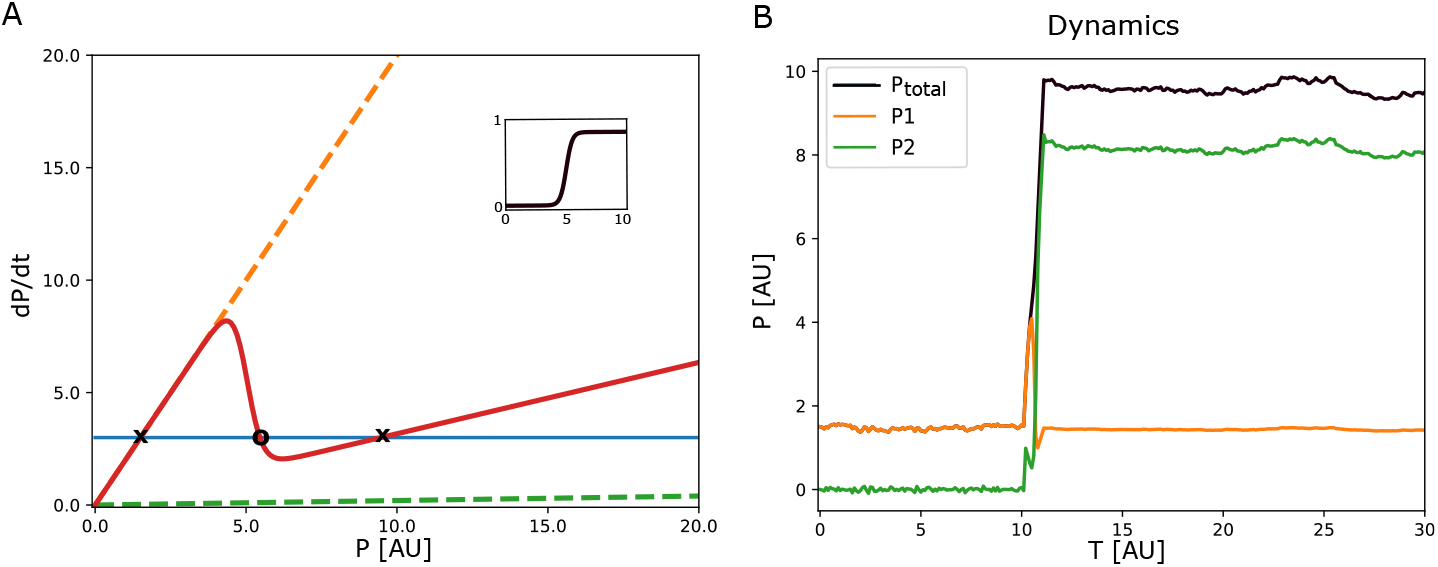
Bistability with negative feedback with a small change in *P*_1_ between DOWN and UP states. **A.** Sink and Source terms when the transition function (*f*) saturates below 1 (inset), specifically *f*_*max*_ = 0.85. The ratio of *λ*_*1*_*/λ*_*2*_ = 100. There are two stable fixed points (x) and one unstable fixed point (o). **B**. Dynamics of the saturating model. Induction from 10 — 11 in the arbitrary time units. The level of *P*_*2*_ increases significantly, but *P*_*1*_ stays relatively stable. Before induction (*T <* 10) the level of *P*_*total*_ = *P*_*1*_ and therefore *P*_*total*_ cannot be seen because it is obscured by *P*_*1*_.

The form of the state function (*f*) determines if the system is mono- or bistable. One could imagine other functional forms of *f* for which the system might be multi-stable. In the extereme case this system can also become a continuous attractor over a limited range and for a very specific state function. A form of the state function that can generate such a continuous attractor can simply be found by setting equation to zero, and solving for f. This produces the equation:

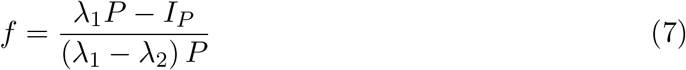

This equation is valid for *P* such that *f* ≥ 0; otherwise it is set to zero.

With this choice of the state-function all levels of P in the range *I*_*p*_*/λ*_1_ ≤ *P* ≤ *I*_*p*_*/λ*_2_ are stable; this is the range of continuous stability. In Figure 4A we show the source and sink terms given this form of the state-function, and in the inset the form of f is displayed. In Figure 4B we show the results of several different consecutive induction episodes triggered by a transient increase in *I*_*P*_. Every transient causes a change in the steady state of the system, as long as *P* stays within the allowed range. This continuous attractor is an integrator as it keeps a stable record of the integral of the inputs into the source term. With no noise as in Figure 4B, these steady states are stable. However, if noise is added to these simulations (Fig. 4C), there are fluctuations of the state within the allowed range.

**Figure 4.**
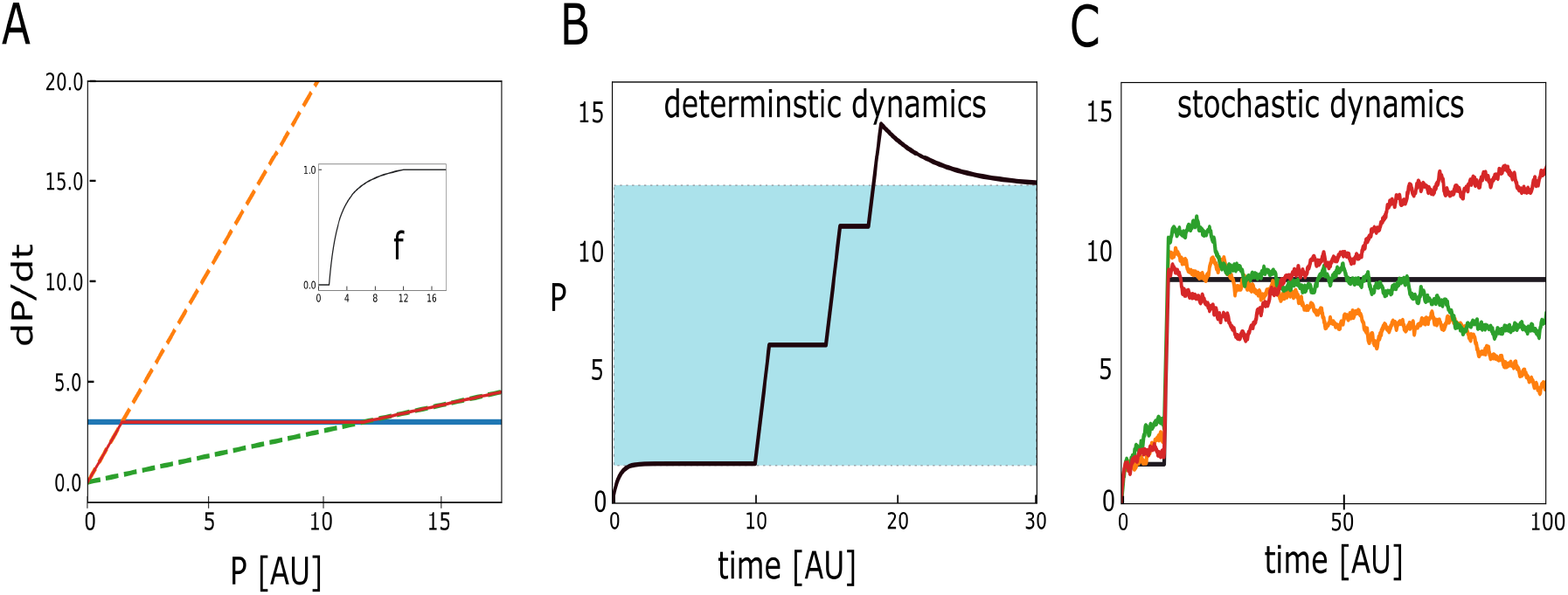
A continuous attractor resulting from well-tuned negative feedback. **A.** The source and sink terms, labeled as in previous figures. The sink and source terms have a substantial overlap by construction, as determined by using the state-function in equation 7 (see inset). **B**. Simulating the continuous attractor model. Initially the model settles at the lowest possible persistent state where *P* = *I*_*P*_ */λ*_*1*_. A set of three consecutive induction stimuli cause jumps to new persistent levels of *P*. The last stimulus causes a large jump which decays back to the maximal allowed persistent level of *I*_*P*_ */λ*_*2*_. The light blue shaded area represents the range of continuous stability. **C**. Stochastic simulations of the system. The deterministic simulation (black) exhibits persistent activation. Three different stochastic simulations (orange, red, green) show significant fluctuations within the “stable” region. In simulations the parameters are: *I*_*P*_ = 3, *λ*_*1*_ = 2, *λ*_*2*_ = 0.25, all in arbitrary units [AU]. The added noise in C has mean 0 and a standard deviation of 0.25 in [AU].

The magnitude of the fluctuations, as well as the velocity of the drift, increase with the noise level. Such fluctuations within the stable range are an inherent property of continuous attractors. Note that the time axis here is labeled with arbitrary units ([AU]), which means that we do not know the time scale of the drift along the continuous attractor. With such a simple and abstract model there is no way to set real time units. With more complex models in which actual molecular reactions such as synthesis, diffusion, degradation and binding take place, it would be possible to set physical time units, but only once these biophysical parameters are estimated.

To summarize, in the abstract model of maintenance based on negative feedback of the sink term, the non-linearity responsible for bistability is in the sink term, rather than in the source term. A necessary condition for obtaining stability is the steep (ultrasensitive) state function. Here we assume it exists but do not explain its mechanistic origin. In section 2.4 we propose a more detailed model motivated by experimental evidence regarding interactions between specific molecules. That model will no longer be a 1D model, though it still belongs to the negative-feedback theoretical framework. Many of the major qualitative features of the abstract model, including the dependence on the different degradation rate constants and the critical importance of the ultrasensitive process, are maintained.

### 2.4 A biophysical model based on KIBRA - PKMζ **interactions**

Recent results have shown that L-LTP and long-term memory maintenance crucially depend of the continual binding between two synaptic proteins, *PKMζ* and KIBRA (Tsokas et al., 2024). After L-LTP the concentration of the bound proteins appears to increase selectively in synaptic spines. Molecules that selectively inhibit the binding of these proteins can reverse L-LTP without affecting basal synaptic transmission. These selective inhibitors can also disrupt memories up to a month after they were established. It is also known that the binding of *PKMζ* to KIBRA shields *PKMζ* from proteasomal degradation, increasing the kinase’s steady-state levels in cells (Vogt-Eisele et al., 2014; Hu et al., 2017; Ferguson et al., 2019; Tsokas et al., 2024).

We have previously shown using proximity ligation assay (PLA) that *PKMζ* and KIBRA bind, and that this binding is essential for memory maintenance (Tsokas et al., 2024). We have also shown, using an expression system composed of HEK293 cells, that the co-expression of *PKMζ* and KIBRA results in small regions, in which there is a high density of both proteins (Tsokas et al., 2024). In supplemental Figure 1A we show, using bimolecular fluorescence complementation (BiFC), that high-density regions of bound KIBRA bound to *PKMζ* are formed. We also show that *ζ*-stat K-ZAP, which prevent the binding of KIBRA and *PKMζ* eliminates these high-density regions (supplementary Figures 1C,D). We show that the formation of these high-density regions requires the existence of KIBRA and *PKMζ* and is not an artifact of BIFC (supplementary Fig. 2).

We have reanalyzed our previously published results of the binding of *PKMζ* and KI-BRA in HEK293T (Tsokas et al., 2024) cells to calculate their steady-state concentrations (Supplementary Figure 1E,F). We found that if their binding is prevented using *ζ*-stat or K-ZAP (Tsokas et al., 2024), their steady-state levels decrease. These results further indicate that the binding of *PKMζ* and KIBRA, which results in the formation of apparent droplets, inhibits the elimination of both of these proteins from the cell.

Although these results are based on BiFC an irreversible reporter of binding, similar results were obtained with PLA which does not suffer from this limitation (Kauwe et al., 2024; Ferguson et al., 2019). In addition, supplementary Figure 2 supports the observation that high-density regions of co-localized KIBRA and *PKMζ* are formed due to their interactions and not due to BiFC, because high-density regions of co-localized KIBRA and*PKMζ* appear in the absence of the Venus construct necessary for BiFC.

These high-density regions of bound proteins are not solid aggregates, and their structure is reversible. We demonstrate this using photobleaching of these aggregates in the HEK293T cells. This photobleaching results in a rapid reduction in signal, and a recovery of about 25% in the same regions, within 180 seconds (Supplementary figure 3). This indicates that these high-density regions are not solid. However the reappearance of these regions in the same locations indicates that while the molecules themselves are transient, the structure is more stable. Together, these results suggest, in part, that these high-density bound structures might be droplets, or aggregates that rapidly exchange molecules with the surrounding low-density regions.

Based on the observations that the binding of KIBRA and *PKMζ* is essential for maintaining L-LTP and memory, we developed a biophysical model of maintenance. The model is also motivated by our observations that bound KIBRA and *PKMζ* produce larger structures, but that these structures are not completely solid as demonstrated by the recovery from photobleaching.

From a theoretical perspective the formation of heterodimers of KIBRA and *PKMζ* is not sufficient for generating a bistable system because this first-order reaction is not ultrasensitive. To examine whether the binding of KIBRA and *PKMζ* is likely to produce larger structures, we used AlphaFold3(Jumper et al., 2021; Abramson et al., 2024) to investigate the possible formation of such structures. We find that various larger complexes of KIBRA and *PKMζ* heterodimers, from tetramers to hexamers, are potentially stable. Our results point to stable bound structures from hetero dimers to Hexamers. On the basis of these findings we propose biophysically plausible maintenance models based on the binding of KIBRA and *PKMζ* and the formation of larger complexes of such bound molecules.

In figure 5A we show a schematic diagram of KIBRA-*PKMζ* complexes which form the framework of our proposed biophysical maintenance model. Several options are suggested by AlphaFold3 for each complex; we have chosen to show the most likely one. Given that such higher-order structures are plausible, we have decided to generate dynamical models that include these higher-order structures. The minimal model that was able to generate bistability was the Hexamer model.

**Figure 5.**
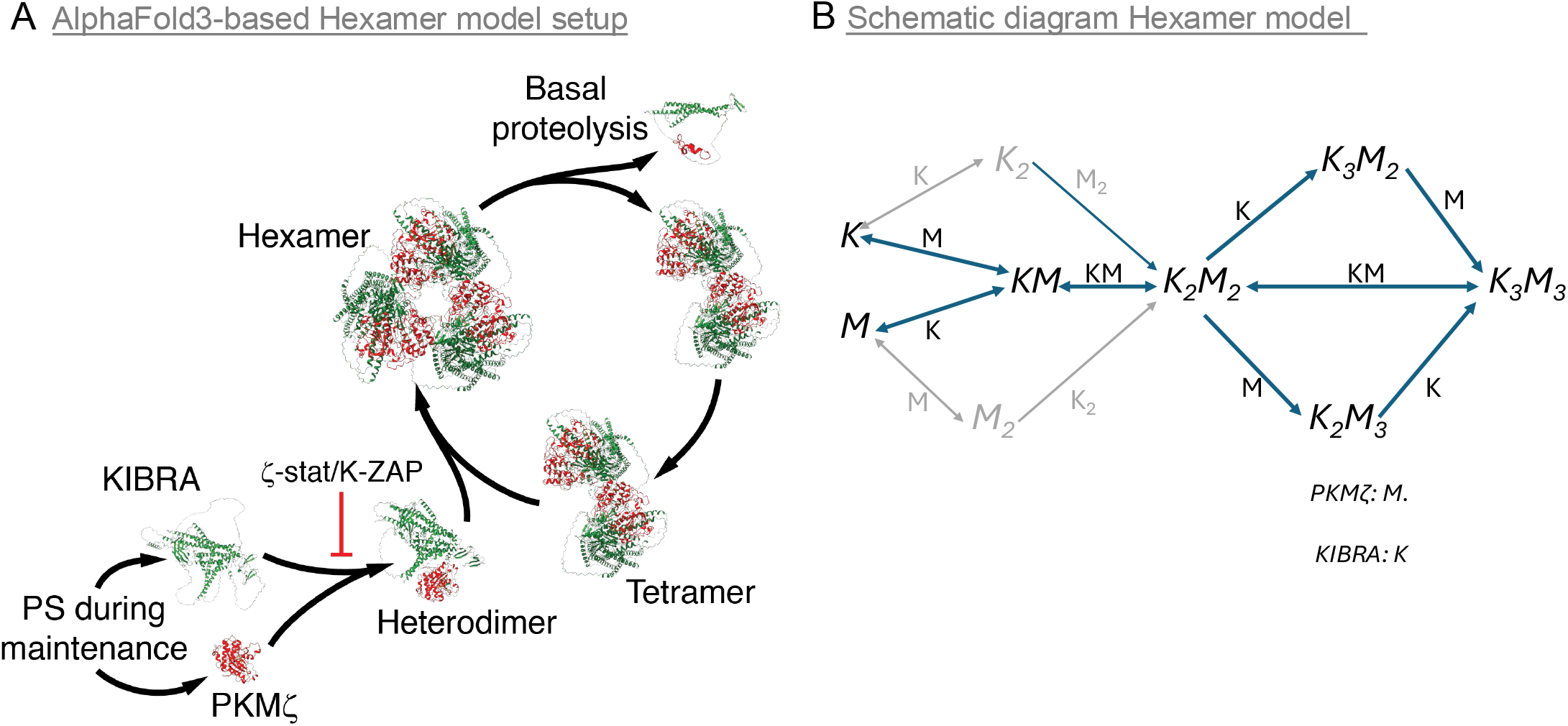
Setup and motivation of the biophysical model based on the binding of KI-BRA and PKM*ζ*. **A.** A simplified diagram of the reactions during maintenance and its disruption by binding inhibitors, based on AlphaFold3 predictions of different of KIBRA-*PKMζ* complexes. KIBRA in green, *PKMζ* in red. We show monomers, dimers, tetramers, and hexamers. The increased amounts of *PKMζ* and KIBRA in the UP state are stable with basal rates of protein synthesis (PS) and turnover (proteolysis shown). **B**. A schematic diagram of all the reactions assumed in the model. There are 7 different species. We use the notation *K* for KIBRA and *M* for *PKMζ*. The subscript numbers denote the number of molecules of each type in the complex. For example, *K*_2_*M*_3_ denotes a complex with two KIBRA and three *PKMζ*. The letters above the arrows indicate the binding partner. In addition to the reactions shown there is protein synthesis of *K* and *M*, and all species can degrade. The degradation of the monomers (K and M) is much faster than of the more complex species. The pathways through *K*_2_ and *M*_2_ are shown in gray as they are not necessary and not used in the results shown in Figure 6, though they can be and are used in the different parameter sets supplied, and in figure 7.

In Figure 5B we show a schematic diagram of the species and reactions included in the model. In addition to the reactions shown, the model includes constant protein synthesis of KIBRA and *PKMζ*, which only increases transiently during the induction phase. It also included degradation, elimination, or diffusion of the different species, where the monomers and dimers are eliminated at a much faster rate than the more complex species. The mathematical details of the model are discussed in Appendix A. Qualitatively, the model is similar to the abstract model described above, where the complex species provide the sufficient non-linearity and the differential degradation are key assumption of the abstract model described in section 2.3.

This model can be bistable as shown in the simulation shown in Figure 6. In Figure 6A we show the dynamics of all species, and in Figure 6B we show the dynamics of the total of KIBRA and *PKMζ*. The model is initialized with low initial conditions and allowed to converge to the DOWN state. Then a brief induction protocol for L-LTP is applied by transiently increasing the protein synthesis rates of KIBRA and *PKMζ*. The model then transitions to the UP state. This input pulse is equivalent to a transient increase in protein synthesis of key synaptic proteins which is essential for the induction of L-LTP and long-term memory in general (Klann and Sweatt, 2008).

**Figure 6.**
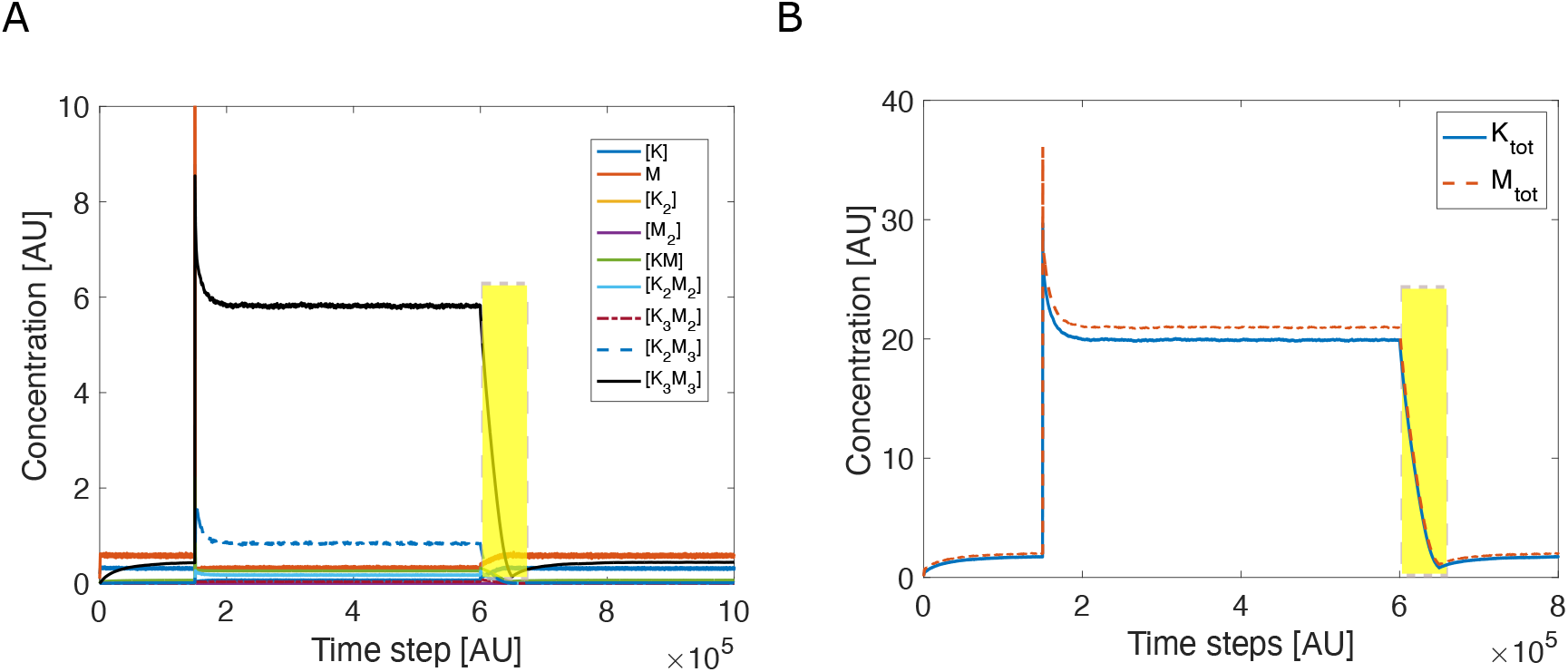
Induction and reversal of L-LTP with the Hexamer model. **A.** Model dynamics of all species in the model, see legend. The induction fo L-LTP at 1.5 *·* 10^5^ time units causes a rapid but sustained increase in the complex species, especially *K*_3_*M*_3_ (black). The transient inhibition of binding between KIBRA and *PKMζ* (yellow rectangle) results in a sustained transition back to the previous state. **B**. The same as in A but for total KIBRA (*K*_*tot*_ - blue) and total *PKMζ* (*M*_*tot*_ - red).

The allosteric inhibitors *ζ*-stat or K-ZAP that act by inhibiting the binding of *PKMζ* to *KIBRA* reverse L-LTP and long-term memory (Tsokas et al., 2024). We simulated the application of *ζ*-stat or K-ZAP by transiently reducing the binding coefficient between *PKMζ* and *KIBRA*. This transient reversal protocol (yellow rectangle) is sufficient to cause the model to transition back to the DOWN state, as shown in Figure 6 (t=6 [AU], yellow bar). In the model, the application of *ζ*-stat or K-ZAP at the basal level causes only a minor and transient change in total protein levels (Results not shown).

L-LTP in the model is induced by a very brief increase in protein synthesis. However, a shorter induction duration or a smaller increase in synthesis rates would result only in a transient increase in concentrations of the complexed molecules. To reverse the UP-state, as shown in Figure 6, a much longer duration of the application of the inhibitors is required. A shorter duration would result in a transient decrease that will bounce back to the UP attractor state.

There are a wide range of parameters for which this model is bistable. We have attached 5 different parameter sets that generate bistability with the code. With these different parameter sets, bistability is observed, but the quantitative values such as concentrations in the UP and DOWN states are very different. Note that, as in the abstract model, the concentration of the less stable species (Monomers) is lower in the UP state than in the DOWN state. This is seen in Figure 6A, by the stable decrease of monomeric species after the induction of L-LTP at time 2 *·* 10^5^ [*AU*]. This decrease, also exhibited in the abstract models, is a necessary condition for negative feedback models, whether abstract of biophysical.

The Hexamer model is the minimal KIBRA-*PKMζ* model for which we observed bistability. It is quite likely that models with larger complexes would also exhibit bistability, we have not examined this. Additionally, although we could not obtain bistabiltiy based on negative feedback with simpler models, we did not prove that it is not possible.

We have shown analytically (equation 3) for the abstract model with a concentration-dependent degradation rate (section 2.3), that the ratio of protein concentrations in the UP and DOWN states are inversely proportional to the ratio of their turnover coefficients. In some sense this must be true in general, since it is the balance between production and degradation that sets steady-state levels, and since in this model the production level is constant, the only possible source of change that determines steady-state levels is the relative degradation rate. Nevertheless, it is instructive to measure this for simulations of the biophysical model as a benchmark that can be compared to experiments in order to test if they indeed fall into the negative feedback framework. We therefore calculated the effective degradation rate coefficient of KIBRA in the UP and DOWN states. The degradation rate coefficient was calculated by measuring the total turnover of KIBRA from each of the species that include molecules of KIBRA and dividing this by the total concentration of KIBRA, in each of those states (see appendix for details). This produces an effective degradation rate coefficient. In the abstract model of section 2.2, this calculation would produce the value of *λ*_1_ in the DOWN state and *λ*_2_ in the UP state. The same calculation exactly can be done for *PKMζ*.

In Figure 7 we plot the inverse of the effective degradation rate coefficient of KIBRA as a function of the total protein in all the KIBRA species. This calculation is repeated for various sets of parameters that exhibit bistability, and both for the UP (+) and DOWN (x) states. As expected, the linear relationship holds. We show the inverse degradation rate constant rather than the degradation rate constant itself because it results in a simple linear-curve. The same synthesis rates were used for all parameter sets presented in this figure. Using different synthesis rates would result in a different linear curve for each synthesis rate, with a different slope for each synthesis rate. Because we are showing inverse degradation, higher synthesis rates produce smaller slopes and lower synthesis rate produce larger slopes (results not shown). Similar results could be shown for *PKMζ*.

**Figure 7.**
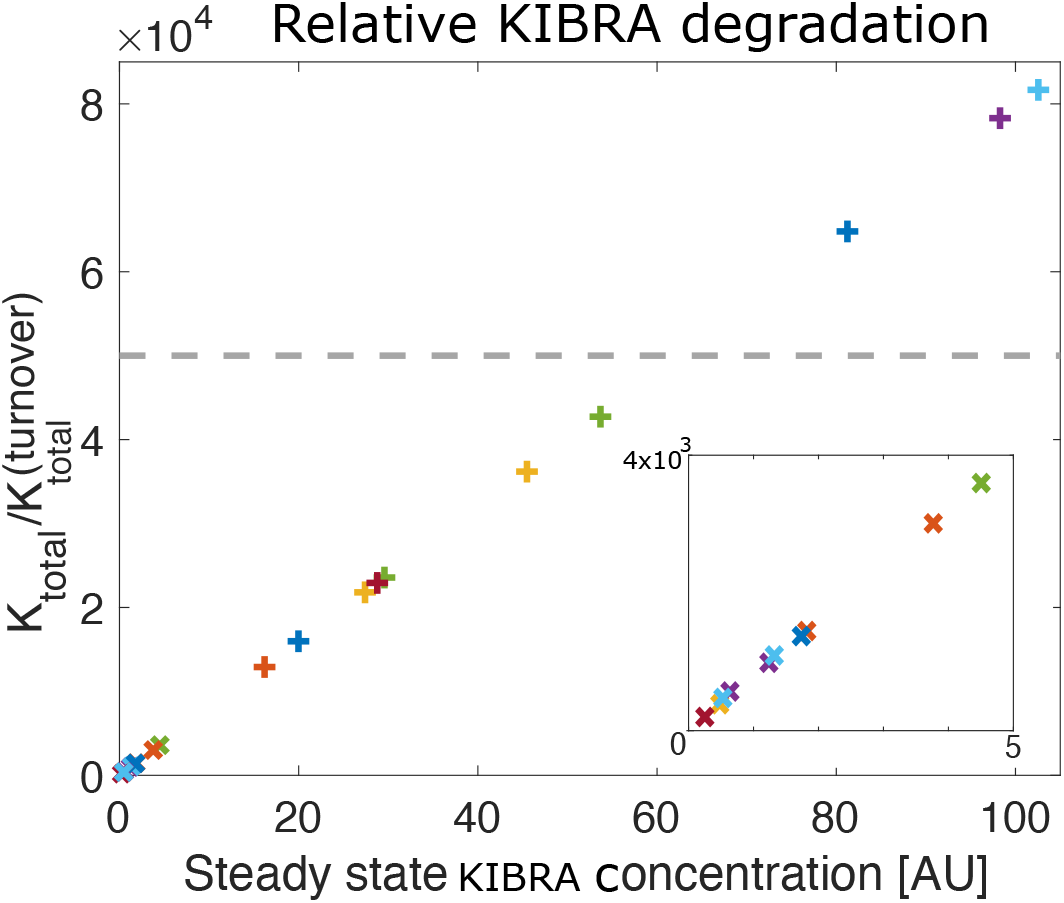
Inverse degradation as function of protein levels in the phenomenological model. Inverse degradation rate coefficients of KIBRA vs. steady state levels of total KIBRA in UP and DOWN states for different parameters. Each color represents a different parameter set, different data points with same color are from the same parameters but for the UP (+ symbols) and DOWN states (x - symbols). All different parameter sets shown have the same synthesis rate. The inset (lower right) shows a blowup of the DOWN state values, which are hard to see in the complete plot due to the very different magnitude of UP state levels for the different parameter sets. The dashed gray line depicts schematically how the inverse degradation rate would behave in a positive-feedback model.

In contrast to this linear relationship exhibited in the negative-feedback model, in a positive-feedback model the inverse degradation rate constant is independent of the state of the system. In the case of the simple positive feedback of section 2.2, the inverse degradation rate constant is simply 1*/λ*, as represented by the dashed gray line of Figure 7.

The inverse effective degradation rate is the time constant for the elimination of molecules in the synapse. In principle, this could be compared to the experimental result of photo-bleaching, as those shown is supplementary figure 7. Currently, the simulation results are in arbitrary units, because the on and off rates used in the model have not been measured experimentally, so a direct comparison is not possible. If we arbitrarily set 10^5^ time steps to an hour (see Figure 6) then the effective time constants for the UP state, shown in figure 7 are between 10 minutes to an hour, on the same order of magnitude as the recovery observed in the photo-bleaching experiment.

The relationship between synaptic efficacy and degradation could be measured experimentally in order to test which of the different theories are consistent with the data. Specifically, the negative-feedback model would predict that after a synapse undergoes L-LTP, the effective degradation rate coefficient of key proteins such as *PKMζ* or KI-BRA would decrease. In a positive-feedback model it would not change. However, these different mechanisms are not necessarily mutually exclusive, and it is possible that both mechanisms are used in the same synapses, i.e., that stronger synapses might exhibit both less degradation and more synthesis. Experimental data could indicate if maintenance is accomplished via a pure positive-feedback or negative-feedback model, or via a hybrid model. If a hybrid model is used, the maintenance system might exhibit multi-stability; enhancing the dynamic range of a synapse.

## 3 Discussion

In this paper we propose a new class of negative-feedback models to account for the stability of synaptic efficacies, and show that such models could account mechanistically for synaptic stability. This class of models are based on two key assumptions. First, in a potentiated state, key synaptic proteins are eliminated at a slower overall rate coefficient relative to the un-potentiated state. Second, there is a steep hypersensitive transition separating the un-potentiated and potentiated states. We show here that this novel framework can qualitatively account for experimental observations of L-LTP and long-lasting memories. We also show that the negative feedback model generates different predictions than the positive feedback models-a difference that can be tested experimentally.

We put forward both an abstract model (section 2.3) and a mechanistic model (section 2.4). The mechanistic model is based on observations that the binding of KIBRA to *PKMζ* is required for the stabilization of synaptic plasticity and memory in wild-type animals (Tsokas et al., 2024). Previous results have shown that the binding of *PKMζ* to KIBRA reduces protein turnover (Vogt-Eisele et al., 2014). In addition we have observed the formation of high-density, droplet-like structures of KIBRA bound to *PKMζ*, and AlphaFlow3 predicts that larger scale complexes of KIBRA bond to *PKMζ* are possible. Thus, we hypothesize that the binding of *PKMζ* to KIBRA can slow elimination of these proteins from synaptic spines, and that the cooperative formation of larger complexes can provide the non-linearity necessary for bistability. Our simulations of many variants of such a model show that a model based on the formation of KIBRA-*PKMζ* hexamers is sufficient to account for synaptic stability. These simulations also show that molecules that inhibiting the binding of KIBRA and *PKMζ* can reverse stable synaptic potentiation and long-term memory. This model is robust as many different parameter sets produce qualitatively similar results.

Our experimental results shown in supplementary figures 1 and 2, and our previous publication (Tsokas et al., 2024) show regions of a high-density of bound KIBRA and *PKMζ*. These regions are droplet-like in their shape. Our results also show that they are not solid, and exchange proteins with the surround (supplementary figure 3). These results suggest liquid-liquid phase separation (Liu et al., 2021; Chen et al., 2020), however, we have not proven this conjecture. Moreover, our biophysical model does not depend on this observation. It is possible that a model based on a phase transition between a diffuse phase and a liquid-liquid separated phase could also account for the stability of synaptic plasticity and memory. This option is not explored here.

The premise that stable synaptic efficacies underlie stable memories and learning has been called the synaptic trace theory (Mongillo et al., 2017). However, recent long-term recording of synapses in vivo using two-photon microscopy has found a large scale turnover of synaptic spines and large changes in their size over a period of several weeks in the CA1 subregion in the hippocampus (Attardo et al., 2015). Experiments that show synaptic volatility seem to contradict the assumption of the synaptic trace theory. In contrast, spines seem to be much more stable in adult sensory and motor cortex (Grutzendler et al., 2002; Zuo et al., 2005). Despite the recent advances in recording techniques we do not yet know if the fraction of stable synapses observed are indeed those crucial for the maintenance of memory.

The apparent instability of synaptic efficacies is mirrored by instability of neuronal representations in some brain systems and conditions (Ziv et al., 2013) but not others (Refaeli et al., 2023; Pérez-Ortega et al., 2021). A drift in neuronal representations might be a consequence of synaptic instability. Models have shown that in some systems a drift of the input representation can be compensated for by upstream mechanisms, and slowed down by external feedback (Rule et al., 2020) or by internal mechanisms (Pérez-Ortega et al., 2021). Such mechanisms require constant re-exposure to the same environment, and this does not apply to memory of old events that one is not re-exposed to. Currently, it is unclear how old memories can be retained if there is no stable synaptic core. Many synapses might still drift, if their drift is in a subspace orthogonal to the stored memories, but currently it seems likely that an episodic memory cannot be maintained without a stable synaptic core (Clopath et al., 2017).

Another possible objection is that a process of system-level consolidation occurs after the induction of long-term memory. In system level consolidation, the location of memory storage is shifted from the hippocampal region to cortical regions (Wang and Morris, 2010; Squire et al., 2015). However, in these cortical regions memory is also stored via synaptic plasticity, which means system level consolidation does not change the nature of the problem, but only its physical location. Despite system level consolidation, it has been shown that engrams in CA1 and neocortex are highly stable for long period of time, consistent with local memory storage (Refaeli et al., 2023; Lee et al., 2023). More directly, we have specifically shown that the reversal of the molecular mechanism maintaining L-LTP in the hippocampus, can disrupt memory storage up to at least one month after the formation of memory (Tsokas et al., 2024). These results show that at least some memories are stored in the hippocampus for at least a month (Tsokas et al., 2024), much longer than typical dwell times of key synaptic proteins.

One must note that models of positive feedback are the prevalent models not only for memory maintenance, but more prominently for models of cell fate (Kobayashi et al., 2003; Zhu et al., 2022). In those systems the feedback loop is typically implemented at the level of transcription. In these theories transcribed proteins either directly or indirectly modulate their own transcription. Models implemented at the level of transcription are whole cell models. Therefore, they are inappropriate for maintenance of synaptic plasticity that must be synapse-specific in order to be able to maintain specific memories, generate selective receptive fields, and for specific learning in general. Although a transcription-based feedback loop is inappropriate as the primary mechanism for maintenance of synaptic plasticity, changes in transcription may also occur during maintenance (Squire et al., 2015).

In order to serve as a basis for memory formation and for learning in general, synaptic plasticity must exhibit a high degree of synaptic specificity. Synaptic specificity means that while some synapses are potentiated, neighboring synapses remain un-potentiated. Such synaptic specificity has been observed experimentally during the induction of LTP (Harvey and Svoboda, 2007) and L-LTP (Govindarajan et al., 2011). Positive-feedback models of maintenance are based on mechanisms that are in principle local such as post-translational modifications, or local translation of proteins. However, until recently the degree of synapse specificity arising from such supposedly local mechanisms has not been analyzed theoretically.

To address the question of synapse specificity we recently developed a reaction diffusion model of maintenance based on positive feedback (Huertas et al., 2022). The model includes a dendrite with many synaptic spines and with molecular switches based on positive feedback. We studied this model using both analytical and numerical methods, and we have shown that in order to obtain synaptic specificity it is necessary that the switches reside in synaptic spines. However, although necessary, this might not be sufficient, as we have also found that even if the bistable switches reside in spines it is very difficult to obtain synaptic specificity at the level observed experimentally. This lack of specificity occurs because proteins generated in active spines can spill over to neighboring inactive spines and turn on the switches in those spines. If a sufficient number of such synapses are in the potentiated state this spillover will cause all their neighbors within the dendritic branch to potentiate as well, resulting in a loss of synaptic specificity. Our results showed that the one way to obtain a realistic level of synaptic selectivity is if in addition to the positive-feedback loop another mechanism during the induction of L-LTP restricts the diffusion of the critical protein out of the potentiated spine. The structural changes that occur in L-LTP (Fukazawa et al., 2003) suggest that such changes in diffusion are possible, but we have shown that changes in diffusion need to be quantitatively quite large, in order to establish realistic levels of synaptic specificity.

It is possible that a negative feedback model will not suffer from the same limits to synaptic specificity. This is because in a negative feedback model, potentiated synapses do not persistently synthesize additional proteins, which then diffuse out of the spines, potentially activating nearby spines. However, we have not yet analyzed the synapse specificity of negative feedback models mathematically.

Stability based on a negative-feedback loop, which is dependent on decreased degradation (presumably by decreasing the energy-consuming ubiquitin-proteosome system), or diffusion might also be more energy efficient than a positive-feedback loop model. Positive feedback models require either a persistent increase in protein synthesis (Aslam et al., 2009; Ogasawara and Kawato, 2010; Jalil et al., 2015; Klann and Sweatt, 2008) or in autophosphorylation (Lisman, 1985; Lisman and Zhabotinsky, 2001; Miller et al., 2005) to maintain the UP state. In contrast the energy requirements for the negative feedback system are similar for both UP and DOWN states. In this family of models no persistent increase in protein synthesis or autophosphorylation, processes that require energy, are necessary to maintain the UP state. In negative feedback models the two stable fixed points are intrinsically stable due to the relative stability of the larger complexes, and the decreased relative protein elimination. Only a transient increase in protein synthesis is required for the induction of L-LTP, consistent with experimental observations (Klann and Sweatt, 2008; Frey and Morris, 1997).

## Acknowledgements

This work is supported by NIH funding RO1 DA034979 (HZS, TCS), RO1 MH53576 (TCS), 2R37 MH057068 (TCS), R01 NS105472 (TCS), R01 NS108190 (TCS), and MH115304 (TCS).

## APPENDIX A The biophysical model based on KIBRA-PKMζ interactions

The Hexamer model is based on elementary reactions between KIBRA and *PKMζ*. We use the abbreviation K for KIBRA and M for *PKMζ*. There are 7 distinct species: *K, M, KM, K*_2_*M*_2_, *K*_3_*M*_2_, *K*_2_*M*_3_, *K*_3_*M*_3_. The complete set of reactions we consider is below:

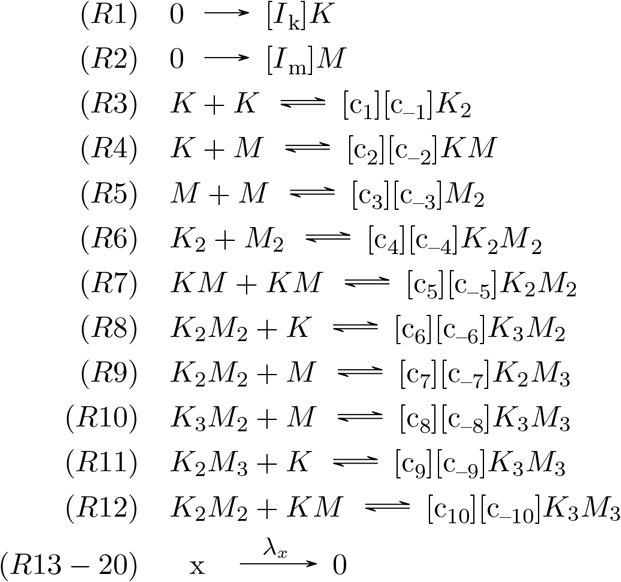

Reactions R13-R20 are for the degradation/elimination/diffusion of proteins from the spine. All proteins (*x*) are eliminated with their own elimination rate (*λ*_*x*_). The main restricting assumptions here are: First, the elimination rates for the monomers are much faster than those for the more complex species, tetramers-hexamers. In addition, there are no trimers to promote cooperative transitions to the more complex species. Under these restrictions, many sets of parameters are possible. In the following, we provide the parameters for the results shown in Figure 6. In the provided code, we give 4 additional distinct sets of parameters that all produce bistability, with very different quantitative values for protein concentrations at the DOWN and UP states.

The parameter values used for Figure 6 are; Reactions on-rates: *c*_1_ = 0, *c*_2_ = 0.002, *c*_3_ = 0, *c*_4_ = 0, *c*_5_ = 0.02, *c*_6_ = 0.1, *c*_7_ = 0.1, *c*_8_ = 0.1, *c*_9_ = 0.1, *c*_10_ = 0.04. Reactions off-rates: *c*_*—*1_ = 0, *c*_*—*2_ = 0.01, *c*_*—*3_ = 0, *c*_*—*4_ = 0, *c*_*—*5_ = 0.00003, *c*_*—*6_ = 0, *c*_*—*7_ = 0, *c*_*—*8_ = 0, *c*_*—*9_ = 0, *c*_*—*10_ = 0.00125. The degradation rates are 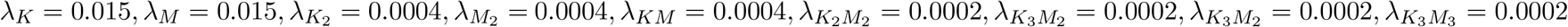. The synthesis rates were: *I*_*k*_ = 0.005, *I*_*m*_ = 0.009. We supplied 4 additional, significantly different, parameter sets that exhibit bistability. For Figure 7 we used 10 different parameter sets (included with the code), all with the same synthesis rates.

These 20 reactions result in 9 coupled differential equations. The simulations were written in Matlab (code provided) using a forward Euler integration method. Noise was added only to the synthesis equations, using a Langevin-like approach to qualitatively test the stability of the fixed points. Stability is determined by running simulations in the presence of noise for a number of time steps that is orders of magnitude larger than the longest time constant in the system. No formal stability analysis is conducted.

For the inhibition we reduce the forward finding coefficients between KIBRA and *PKMζ* by a factor of 2 for 0.5 * 10^5^ arbitrary time units. If each 10^5^ [AU] is one hour, then the inhibition time is 30 minutes. Significantly shorter duration do not reverse the UP state.

To generate Figure 7 we need to calculate the effective degradation coefficients. To calculate the effective degradation coefficient of KIBRA and *PKMζ*, we first calculated the total degradation of *PKMζ* at each time step (*dt*). For this model, the degradation of KIBRA at time t is: 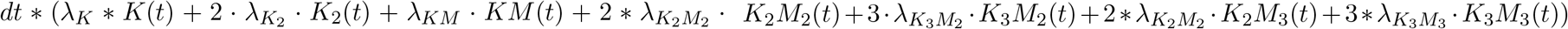. The equivalent expression can be used for *PKMζ*. To obtain the effective degradation rate coefficient we divided the total degradation of KIBRA at each time point by the total concentration of KIBRA at the same time point. These were averaged separately over time point windows in the DOWN and UP states, long after the transitions. The y axis values of Figure 7 are the inverse degradation rates, which are 1 over the effective degradation coefficient. Although noise is added to the synthesis rates, it has little effect on the average values of protein concentrations and degradation levels. An identical procedure could be carried out of *PKMζ* yielding similar results.

## Supplementary Figures

**Supplementary Figure 1.**
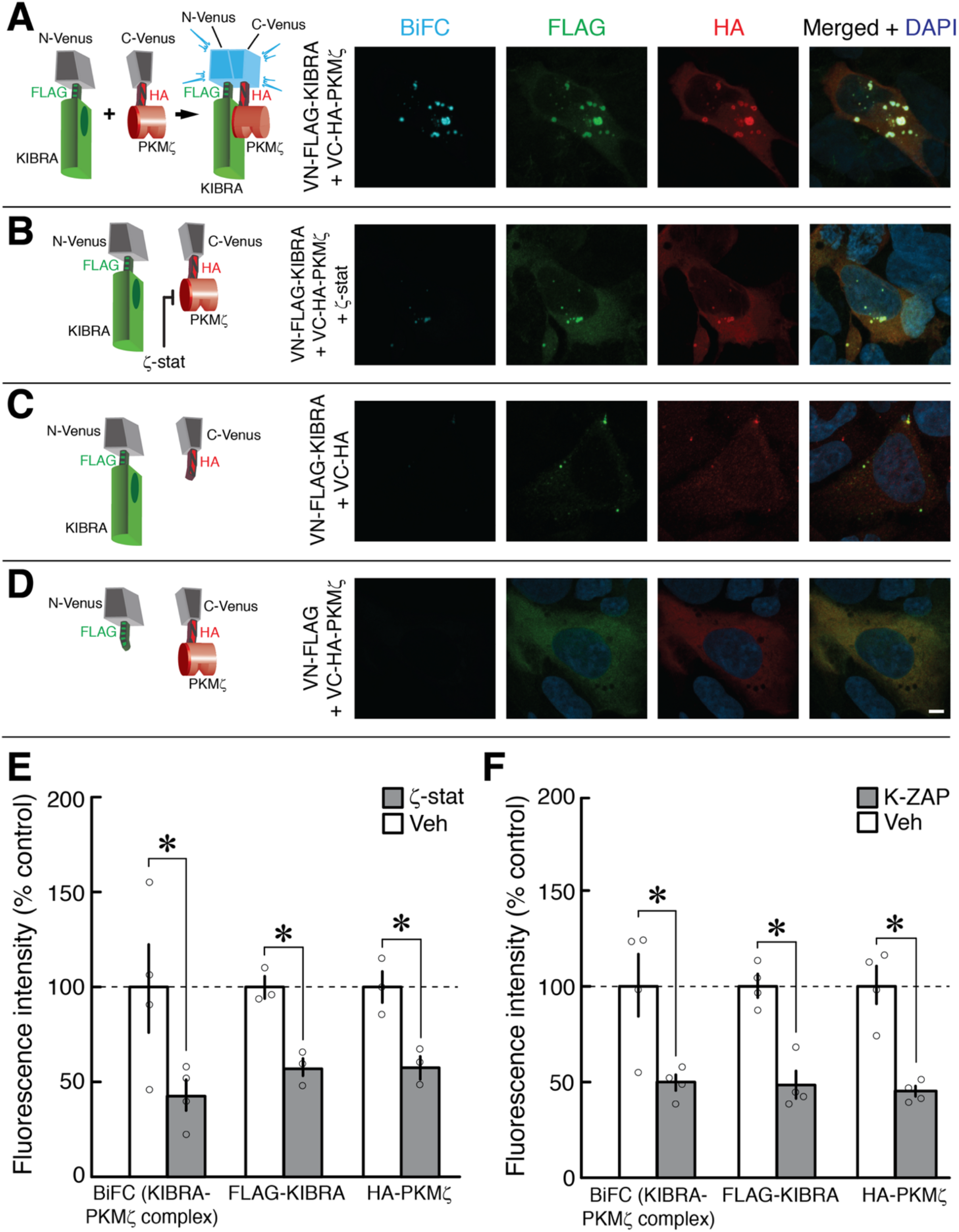
KIBRA-PKMζ antagonists that decrease KIBRA-PKMζ interaction lower total KIBRA and PKMζ levels. **(A-D)** Left, schematics of Venus-fusion constructs used in BiFC experiments. Right, representative images of HEK293T cells 1 day after transfection. Left column, BiFC (cyan); middle columns, immunocytochemistry for FLAG-tagged KIBRA (green) and HA-tagged PKMζ (red); right column, merged images with nuclear stain DAPI (blue). **(A)** BiFC produced by co-transfection of N-terminal-Venus FLAG-tagged KIBRA and C-terminal-Venus HA-tagged PKMζ shows accumulation of intracellular aggregates of KIBRA-PKMζ complexes. **(B)** ζ-stat (10 μM) decreases BIFC as well as total levels of KIBRA and PKMζ. **(C)** Co-transfection of the N-terminal-Venus FLAG-tagged KIBRA and the C-terminal-Venus tagged with HA but without PKMζ shows minimal BiFC. **(D)** Co-transfection of C-terminal-Venus HA-tagged PKMζ and N-terminal-Venus tagged with FLAG but without KIBRA shows minimal BiFC. Scale bar, 5 µm. **(E, F)**, mean of total immunofluorescence intensities per field of BiFC, FLAG, and HA ± SEM shows (E) ζ-stat and (F) K-ZAP (10 μ M) decrease BiFC, total KIBRA, and total PKMζ immunofluorescence intensities (E: *t*_3_ = 3.39, *P* = 0.04, *d* = 1.69; *t*_2_ = 10.09, *P* = 0.010, *d* = 5.82; *t*_2_ = 12.69, *P* = 0.006, *d* = 7.33; F: *t*_3_ = 4.00, *P* = 0.03, *d* = 2.00; *t*_3_ = 6.24, *P* = 0.008, *d* = 3.12; *t*_3_ = 6.08, *P* = 0.009, *d* = 3.04, respectively). * denotes *P* < 0.05. Experiments described in Tsokas *et al*., *Science Adv*., 10, eadl0030 (2024).

### Method

To measure fluorescence intensity of BiFC, FLAG-KIBRA, and HA-PKMζ, the images were converted to grayscale using ImageJ 1.52n. Individual cells were outlined using freehand or wand selection tools and added to the region of interest manager. Fluorescence intensity was extracted from each channel for individual cells using the Multi Measure algorithm.

**Supplementary Figure 2.**
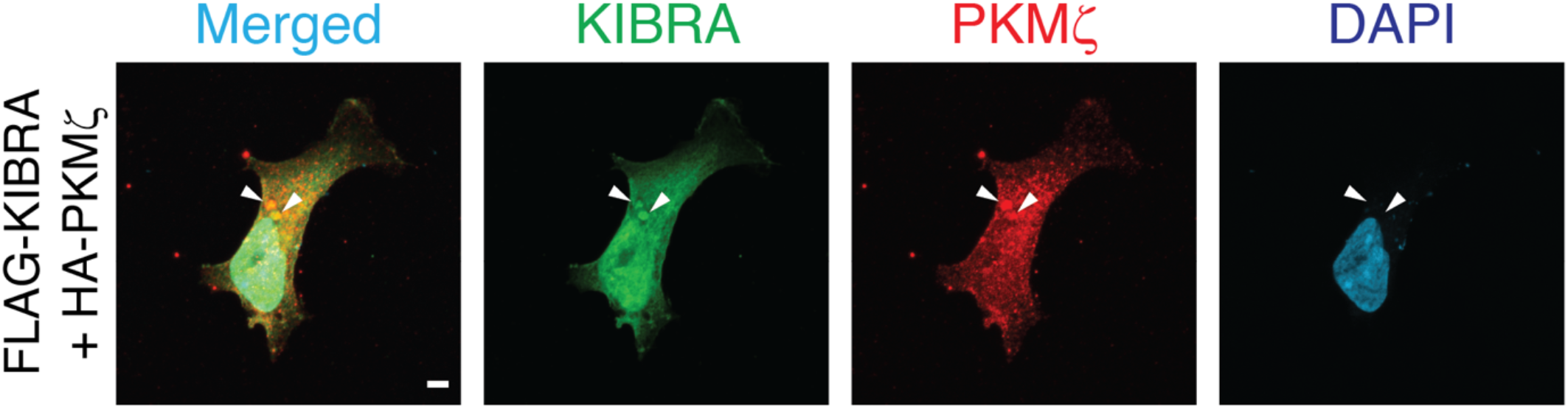
Co-transfection of FLAG-tagged KIBRA and HA-tagged PKMζ without N-terminal or C-terminal Venus shows co-localization in aggregates. Transfections in HEK293T cells were performed as described in Tsokas *et al*., *Science Adv*., 10, eadl0030 (2024). Aggregates are shown at arrowheads. Scale bar, 5 µm.

**Supplementary Figure 3.**
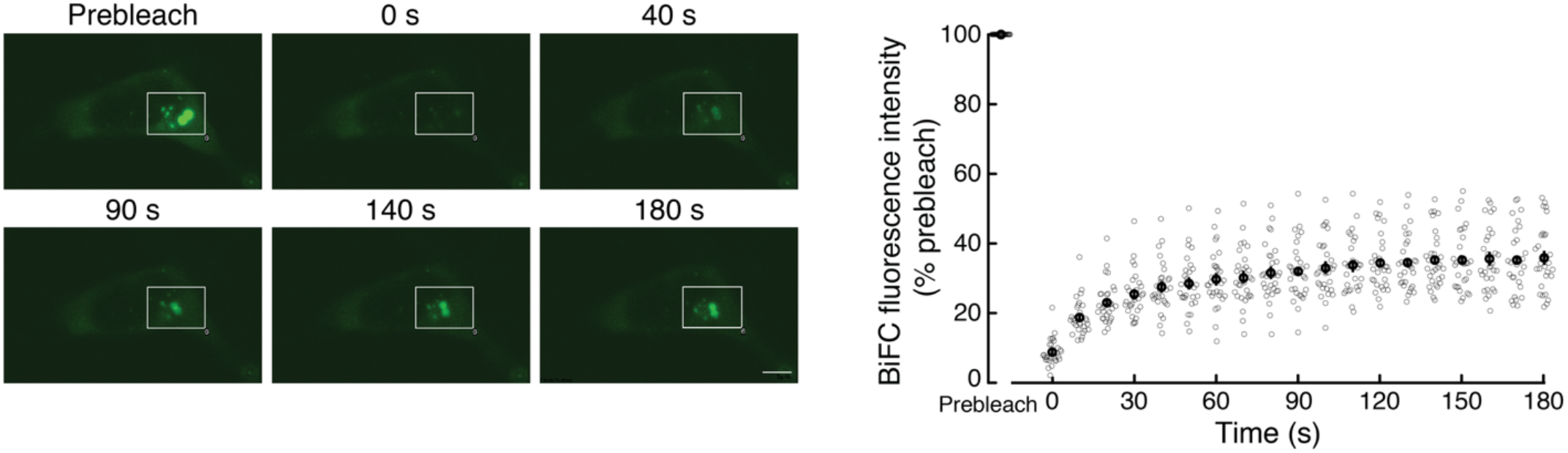
Fluorescence recovery after photobleaching (FRAP) of BiFC shows aggregates contain a mobile pool of KIBRA-PKMζ complexes. The mobile fraction is detected by recovered fluorescence measured within a photobleached region of interest (white rectangle). Left, representative experiment, with postbleaching times denoted. Right, mean (black open circles) ± SEM; n = 30 (10 cells from each of 3 independent HEK293T cultures; individual experimental data shown in grey open circles). Compared to prebleach, the signal is significantly different using Bonferroni correction for multiple comparisons at the first time point postbleaching (prebleach vs. 0 s, paired *t*_29_ = 136.60, *P* < 0.000001), and the signal at 180 s is significantly increased compared to the first postbleaching time point (0 s vs. 180 s, paired *t*_29_ = 18.54, *P* < 0.000001).

### Method

FRAP was performed using an Olympus FluoView FV1000 inverted confocal microscope equipped with a UPLSAPO 60X (NA 1.35) objective (Olympus, Tokyo, Japan) and a stage-top incubator. HEK 293T cells were co-transfected with N-terminal-Venus FLAG-tagged KIBRA and C-terminal-Venus HA-tagged PKMζ, as previously described (Vogt-Eisele *et al*., *J. Neurochem*., 128:686 [2013]; Tsokas *et al*., *Science Adv*., 10, eadl0030 [2024]) and maintained at 37°C in a humidified atmosphere with 5% CO_2_. A region of interest with KIBRA-PKMζ BiFC aggregates was photobleached using a 405-nm SIM Tornado laser (Olympus) at 60% power for 500 ms. Fluorescence recovery in the region of interest was monitored at 488 nm by acquiring images at time intervals denoted in the figure. Image acquisition was performed using Olympus FluoView software, and analysis was conducted with ImageJ software (NIH). Fluorescence intensity of the ROI from each time interval was extracted using the Series Analysis Timelapse algorithm from Olympus FluoView software, and further analysis was conducted with ImageJ software (NIH). Fluorescence intensity of the photobleached area at each time point was background-corrected and expressed as the change in intensity relative to the prebleach intensity set at 100%. Because photobleaching includes areas outside the aggregates, the horizontal asymptote is likely an underestimate of the mobile fraction.

